# Regulation of ROP GTPase cycling between active/inactive states is essential for vegetative organogenesis in *Marchantia polymorpha*

**DOI:** 10.1101/2024.04.05.588222

**Authors:** Yuuki Sakai, Aki Ueno, Hiroki Yonetsuka, Tatsuaki Goh, Hirotaka Kato, Yuki Kondo, Hidehiro Fukaki, Kimitsune Ishizaki

**Affiliations:** Graduate School of Science, Kobe University, Kobe, Japan; Graduate School of Science and Technology, Nara Institute of Science and Technology (NAIST), Ikoma, Japan; Graduate School of Science and Engineering, Ehime University, Matsuyama, Japan; Graduate School of Science, Osaka University, Toyonaka, Japan

**Keywords:** Rho/Rac GTPase signaling, bryophytes, organogenesis, cell polarity, land plants ABSTRACT

## Abstract

Rho/Rac of plant (ROP) GTPases are a plant-specific subfamily of Rho small GTP-binding proteins that function as molecular switches by being converted to the active state by guanine nucleotide exchange factors (GEFs) and to the inactive state by GTPase-activating proteins (GAPs). The bryophyte *Marchantia polymorpha* contains single-copy genes encoding ROP (MpROP), two types of GEFs (ROPGEF and SPIKE (SPK)), and two types of GAPs (ROPGAP and ROP enhancer (REN)). MpROP regulates the development of various organs, including the air chambers, rhizoids, and clonal propagule gemmae. While the sole PRONE-type ROPGEF, KARAPPO (MpKAR), plays an essential role in gemma initiation, little is known about the *in-planta* functions of other ROP regulatory factors in *M. polymorpha*. In this study, we focused on the functions of two types of GAPs: MpROPGAP and MpREN. Loss-of-function Mp*ren^ge^* single mutants showed pleiotropic defects in thallus growth, air chamber formation, rhizoid tip growth, and gemma development, whereas MpROPGAP mutants showed no detectable abnormalities. Despite the distinctive domain structures of MpROPGAP and MpREN, Mp*ropgap^ge^*Mp*ren^ge^* double mutants showed more severe phenotypes than the Mp*ren^ge^* single mutants, suggesting redundant functions of MpROPGAP and MpREN in gametophyte organogenesis. Interestingly, overexpression of Mp*ROPGAP*, Mp*REN*, and *dominant-negative* Mp*ROP* (Mp*ROP^DN^*) resulted in similar air chamber defects, as well as loss-of-function of Mp*REN* and Mp*ROPGAP* and overexpression of *constitutively active* Mp*ROP* (Mp*ROP^CA^*), suggesting importance of activation/inactivation cycling (or balancing) of MpROP. Furthermore, we proved the contributions of the sole DOCK family GEF, MpSPK, to MpROP-regulated air chamber formation. In summary, our results demonstrate a significant role of the two GAPs in the development of various organs and that the two GEFs are responsible for organogenesis through the control of the MpROP active/inactive cycle in the vegetative growth of *M. polymorpha*.

## INTRODUCTION

Small GTPases serve as molecular switches cycling between a GTP-bound active state and a GDP-bound inactive state in response to upstream signals, thereby regulating various intracellular processes. Activation is facilitated by guanine nucleotide exchange factors (GEFs), which catalyze the dissociation of GDP from small GTPases, allowing a GTP molecule to bind to the space instead of GDP. Once activated, the small GTPases interact with various effectors to transmit downstream signals. The intrinsic GTPase activity of small GTPases hydrolyzes GTP, leading to a GDP-bound inactive state. The hydrolytic activity of small GTPases is generally low, and GTPase-activating proteins (GAPs) enhance GTP hydrolysis. Guanine nucleotide dissociation inhibitors (GDIs) further inhibit activation by interacting with inactive forms of small GTPases. Among the small GTPases, Rho family GTPases are conserved in a wide range of eukaryotes and control cell morphology and movement by regulating the cytoskeleton in yeast and animals (Schwartz 2004).

Plants harbor a unique Rho family of small GTPases known as the Rho of plants (ROPs). ROPs are conserved across all green plant lineages, including green algae and mosses such as *Physcomitrium patens*, monocots, and dicots (Christensen et al. 2003; Elias 2008; Eklund et al. 2010). ROPs play crucial roles in regulating cell polarity in various tissues and developmental processes. In the moss *Physcomitrium patens*, ROPs regulate cell shape, tip growth, and asymmetric division in the filamentous protonema, and also the three-dimensional development of the gametophore tissues (Ito et al. 2014; Burkart et al. 2015; Bascom et al. 2019; Yi and Goshima 2020). In angiosperms, ROPs control various cellular morphological processes, such as the formation of crenulated leaf epidermal cell shapes and the polar growth of root hairs and pollen tubes, by regulating the cytoskeleton and vesicular trafficking (Craddock et al. 2012; Lin et al. 2015; Feiguelman et al. 2018). The DOCK family RhoGEF protein SPIKE (SPK), a homolog of DOCK180 (180 kDa protein downstream of CRK), and plant-specific PRONE-type ROPGEF activate ROP GTPases by promoting the exchange of GDP to GTP (Feiguelman et al. 2018). Additionally, two classes of ROP GTPase-activating proteins (GAPs), Cdc42/Rac-interactive binding (CRIB) motif-containing ROPGAPs and Pleckstrin homology (PH) domain-containing GTPase-activating proteins (PHGAPs), have been identified (Wu et al. 2000; Eklund et al. 2010; Feiguelman et al. 2018).

Research on the physiological processes regulated by ROP activation and inactivation has advanced using the model angiosperm, *Arabidopsis thaliana*. RhoGEFs promote ROP activation and membrane accumulation (Ou and Yi 2022). The sole DOCK family protein in *A. thaliana*, AtSPK1, has been shown to control actin cytoskeleton assembly by interacting with the WAVE-ARP2/3 complex and to affect microtubule organization, thereby regulating cellular morphogenesis and cell proliferation pattern (Basu et al. 2008; Yanagisawa et al. 2018; Roszak et al. 2021). PRONE-type AtROPGEFs are phosphorylated and activated by membrane-associated receptor-like kinases (RLKs), which regulate the polarization of ROP activation (Zhang and McCormick 2007; Chang et al. 2013; Denninger et al. 2019). In contrast, the intracellular localization of ROPGAPs and RENs (PH-GAPs) has emerged as a critical factor in the localized activation/inactivation control of ROPs (Feiguelman et al. 2018). For example, AtROPGAPs are localized to the subapical region, thereby spatiotemporally limiting the active ROP domain within the apical part of the tip-growing cell, such as the root hair and pollen tube (Ou and Yi 2022). AtREN2 and AtREN3, which are localized in the preprophase band (PPB) region, inactivate ROP, influence the cytoskeletal organization, and control the orientation of cell division in the root meristem and early embryo (Stöckle et al. 2016). However, *Arabidopsis* has 14 ROPGEFs, a single SPIKE, five ROPGAPs, three RENs, and three ROPGDIs, in addition to 11 ROPs, resulting in high functional redundancy among these regulatory factors, making it challenging to decipher the molecular mechanisms regulating ROP signaling.

*Marchantia polymorpha*, a member of the bryophyte family, serves as an ideal model for understanding ROP signaling. The genome of *M. polymorpha* encodes a ROP, a PRONE-type ROPGEF, a DOCK family RhoGEF SPIKE, a ROPGAP, and a REN (PH-GAP). Recently, MpROP was demonstrated to be a key regulator of various developmental processes, including tip growth in rhizoid formation, cell division pattern control in gemma formation, and air chamber development (Rong et al. 2022; Mulvey and Dolan 2023). Furthermore, the PRONE-type MpROP activator MpROPGEF (KARAPPO) is essential for the initiation of gemma development (Hiwatashi et al. 2019). However, the functions of the MpROP inactivators MpROPGAP and MpREN remain elusive, although screening for mutants with abnormal rhizoid morphology has identified Mp*REN* mutants (Honkanen et al. 2016).

In this study, we established that MpREN plays a pivotal role in air chamber development, rhizoid elongation, and gemma formation, all of which are regulated by MpROP. Our results demonstrate the redundant functions of MpROPGAP in conjunction with MpREN during these organogenetic processes. Furthermore, we unveiled the crucial importance of finely tuning the activation/inactivation cycling of MpROP, that is, inactivation by MpREN (and MpROPGAP) and activation by MpSPK, for the morphogenesis of the air chamber in *M. polymorpha*.

## Results

### RhoGAPs in M. polymorpha

A BLASTP search was performed on the *Arabidopsis* AtROPGAP1 (AT5G22400) and AtREN1 (AT4G24580) amino acid sequences as queries in the Genome Database for *M. polymorpha* (http://marchantia.info/; Montgomery et al. 2020). The homology (E-value < 1e-50) was determined to be high, and a single ROPGAP homolog (Mp6g11120, MpROPGAP) and a single REN homolog (Mp8g09680, MpREN) were identified, respectively. We performed comparative analyses of the amino acid sequences of *M. polymorpha*, *A. thaliana*, and *P. patens* (Figures S1A and S2). Analysis of the amino acid sequence of MpROPGAP with those of AtROPGAP1 (AT5G22400), AtROPGAP2 (AT4G03100), and PpROPGAP3 (Pp3c13_4010) revealed the presence of a CRIB domain and a typical GAP domain in MpROPGAP (Figure S1A). Comparison of the amino acid sequences of MpREN with those of AtREN1 (AT4G24580), AtREN2 (AT5G12150), and PpREN1 (Pp3c9_17460) revealed that the PH, typical GAP, and CC (coiled-coil) domains were highly conserved in MpREN (Figure S2A). Molecular phylogenetic analysis was performed using the amino acid sequences of MpROPGAP and MpREN in *M. polymorpha* and ROPGAPs and RENs in major plant lineages for which whole-genome sequences were available (Figures S1B and S2B). The analysis revealed that MpROPGAP and MpREN belong to the same clade as the homologs of moss, hornwort, and lycophytes, respectively.

### Mp*ROPGAP* and Mp*REN* are expressed in the whole plant body throughout their life cycle in *M. polymorpha*

To elucidate the spatiotemporal functions of Mp*ROPGAP* and Mp*REN* throughout the lifecycle of *M. polymorpha*, we investigated their promoter activities. We isolated the 3,837 bp upstream genomic region of Mp*ROPGAP* (*_pro_*Mp*ROPGAP*) and the 5,045 bp upstream genomic region of Mp*REN* (*_pro_*Mp*REN*). Transgenic plants expressing the GUS reporter under these two promoters showed similar GUS signal patterns, with prominent activity in the meristem notches and dorsal air chambers in one-week-old gemmalings (Figure 1A, B). In three-week-old thalli, high promoter activity extended over the thalli, particularly in the meristem notches and midribs (Figures 1C, D). Within the gemma cups, stronger signals were observed on the base floors and in developing gemmae (Figures 1E, F). Although the GUS signals were low in mature gemmae, prolonged staining revealed higher signals around the meristem notches and in certain rhizoid precursor cells (Figure S3A). Rhizoids displayed notable promoter activity in both transgenic lines (Figures 1G, H). These results were consistent with those of the transcriptome and RT-qPCR analyses. Analysis of transcriptomic information from open sources in the MarpolBase Expression database (http://dev-marchantia.annotation.jp/mbex/; Kawamura et al. 2022) revealed sustained expression of both Mp*ROPGAP* and Mp*REN* throughout the lifecycle of *M. polymorpha* (Figures S3B, C). RT-qPCR was performed using RNA extracted from various tissues of *M. polymorpha*. Mp*ROPGAP* was expressed at higher levels in the reproductive organs than in other tissues, whereas Mp*REN* was ubiquitously expressed in all tissues (Figure 1I, J). These results suggest that Mp*ROPGAP* and Mp*REN* are expressed in various tissues and organs throughout the life cycle of *M. polymorpha*.

**Figure 1.**
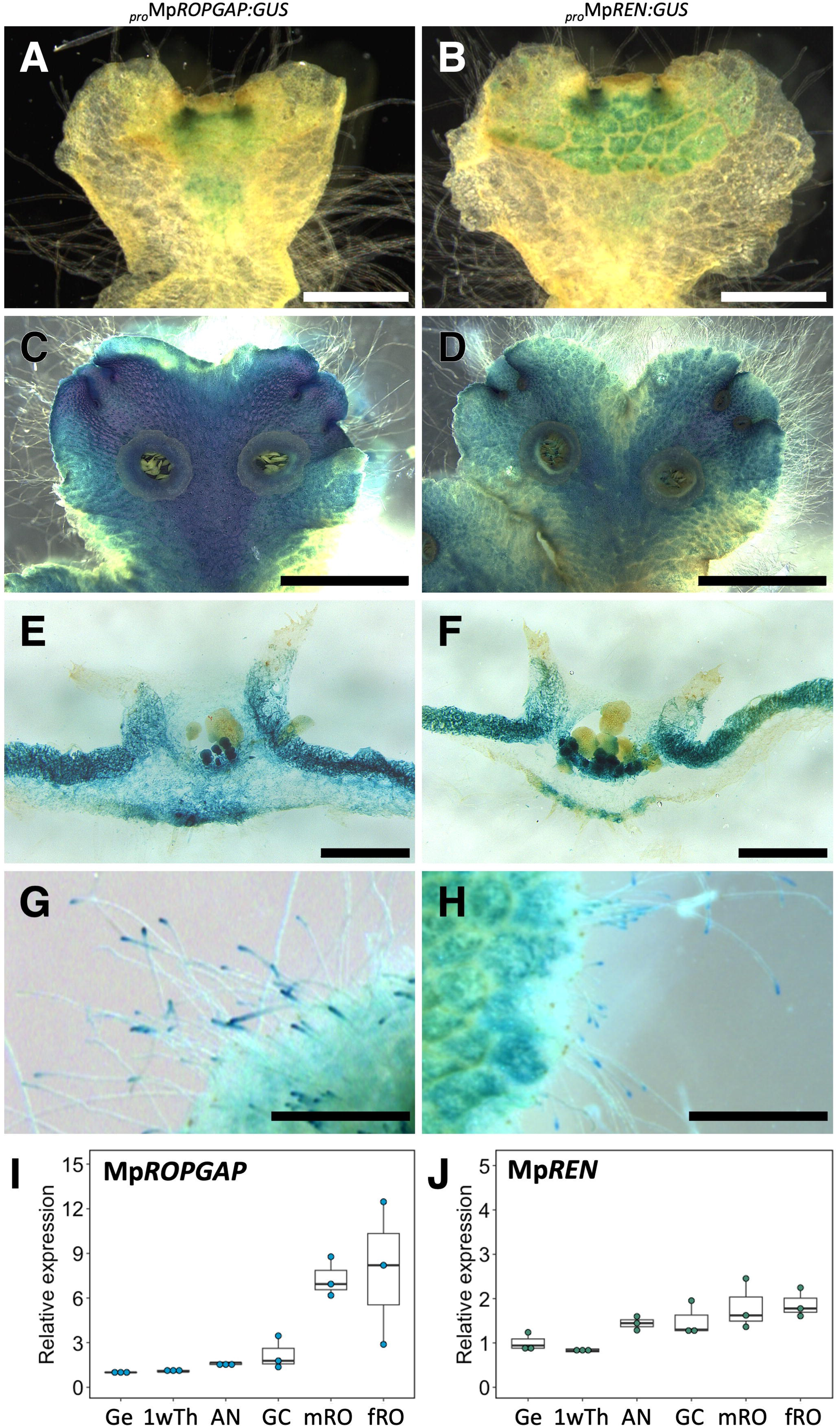
Expression of Mp*ROPGAP* and Mp*REN* in *M. polymorpha*. **(A–H)** Histological GUS staining in representative transgenic lines of *_pro_*Mp*ROPGAP:GUS* (A, C, E, and G) and *_pro_*Mp*REN:GUS* (B, D, F, and H). One-week-old gemmalings (A, B), 3-week-old thalli (C, D), transverse sections of gemma cups on 3-week-old thalli (E, F), and rhizoids on 2-week-old gemmalings (G, H). Bars = 1 mm in A, B, and E–H. Bars = 5 mm in C and D. **(I and J)** qRT-PCR analysis of Mp*ROPGAP* and Mp*REN* in mature gemmae in gemma cups (Ge), 1-week-old thalli (1wTh), apical notch (AN), and gemma cups containing developing gemmae (GC) on 3-week-old thalli, male and female reproductive organs (mRO, fRO). Mp*EF1*_Cl_ was used as the control gene. n = 3.

### MpREN and MpROPGAP are essential for thallus growth and rhizoid elongation

To investigate the functions of ROPGAP and REN in *M. polymorpha*, loss-of-function mutants were generated using the CRISPR-Cas9 system (Sugano et al. 2018). Guide RNAs (gRNA1 and gRNA2) targeting the exon regions encoding the CRIB or GAP domains of MpROPGAP were designed, and genome-edited lines were isolated (Figure S4A). Similarly, guide RNAs (gRNA3–gRNA5) targeting the exon regions encoding the GAP or PH domains of MpREN were designed, and genome-edited lines were isolated (Figure S4B). In these genome-edited plants, frameshift mutations lead to the appearance of early stop codons in the expected transcript sequences. The predicted amino acid sequences lacked the essential domain structures for the functions of MpROPGAP and MpREN, suggesting that these genome-edited strains are loss-of-function mutants (Figure S4C). Furthermore, by conducting genome editing of the Mp*REN* gene in the Mp*ropgap^ge^* -1 mutant background, several Mp*ropgap^ge^* Mpp*ren^ge^*double mutant lines were generated (Figures S4B, C).

Mp*ropgap^ge^* single mutants exhibited a phenotype similar to that of the wild-type plants, displaying normal thallus growth and organ formation, including gemma cups (Figures 2A, B). In contrast, the Mp*ren^ge^* single mutants showed a smaller thallus, and the formation of gemma cups was significantly reduced (Figure 2C). In the Mp*ropgap^ge^* Mp*ren^ge^* double mutants, the thallus phenotypes were more severe than those in the Mp*ren^ge^*single mutants, and no gemma cups were formed during the one-month culture (Figure 2D). We wondered whether the short rhizoids in Mp*ropgap^ge^* Mp*ren^ge^*double mutants restricted access to water and nutrients leading to systemic growth inhibition in the mutant, resulting in the absence of gemma cups. Therefore, we examined the effect of reducing the solidity of the growth medium and increasing the contact between the thallus and the medium. The severe dwarf phenotype of the double mutant was alleviated on media with a low agar concentration of 0.5%, resulting in the formation of distorted gemma cups (Figure 2D, arrow). Distorted gemma cups on the Mp*ren^ge^* single and Mp*ropgap^ge^* Mp*ren^ge^* double mutants generated a small amount of gemmae. These findings suggest that the Mp*ropgap^ge^*, Mp*ren^ge^*, and Mp*ropgap^ge^* Mp*ren^ge^*mutants possess the ability to form gemma cups producing gemmae. Gemma size was comparable to that of the wild type in Mp*ropgap^ge^* single mutants, smaller in Mp*ren^ge^* mutants, and further reduced in Mp*ropgap^ge^* Mp*ren^ge^* double mutants (Figure 2U). The growth of gemmalings during the 6 days from gemmae was comparable to that of the wild type in Mp*ropgap^ge^* single mutants but reduced significantly in the Mp*ren^ge^* mutant and further decreased in Mp*ropgap^ge^* Mp*ren^ge^* double mutants (Figure 2V). In the Mp*ren^ge^* single mutant, abnormal upward growth of the thalli was observed (Figure 2K). To address the potential size evaluation bias due to upward growth, we assessed the flesh weight of plants after 15 days of cultivation, confirming similar results (Figure 2W). Additionally, whereas wild-type plants exhibited bidirectional growth from the gemma, the Mp*ren^ge^* mutant often grew in three or four directions (Figures 2C, G). The growth direction of the Mp*ropgap^ge^* Mp*ren^ge^* double mutant was diverse, with single or multiple directions (Figures 2D, H).

**Figure 2.**
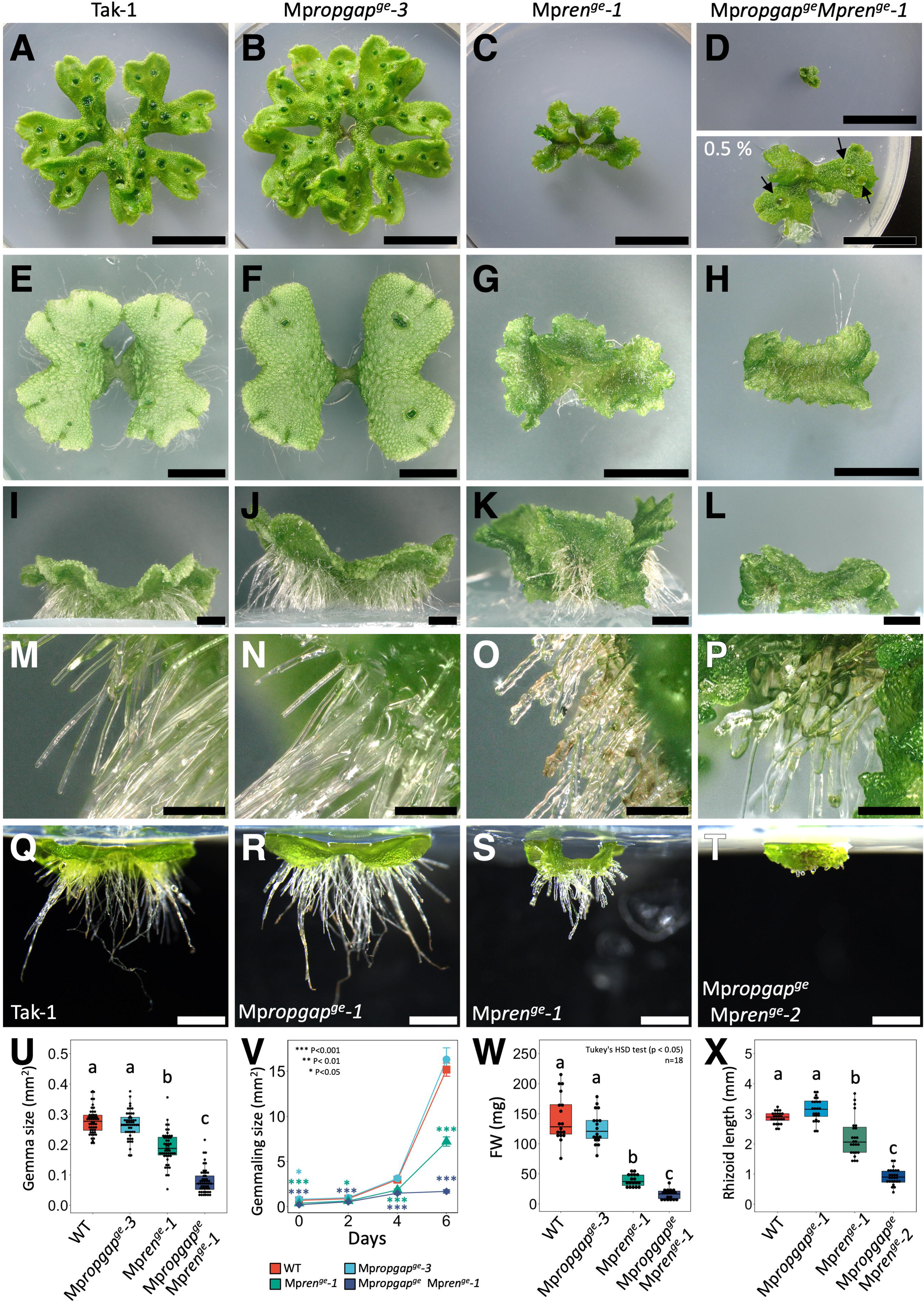
Characterization of Mp*ROPGAP* and Mp*REN* loss-of-function mutants. **(A–P)** Four genotypes are presented, one in each column: wild type (first column; A, E, I, and M), Mp*ropgap^ge^-3* (second column; B, F, J, and N), Mp*ren^ge^-1* (third column; C, G, K, and O), and Mp*ropgap^ge^*Mp*ren^ge^-1* (fourth column; D, H, L, and P). (A–D) Top view of a 4-week-old thalli grown from gemmae. The image below in D is Mp*ropgap^ge^*Mp*ren^ge^-1* grown on a medium containing a lower concentration of agar (0.5%). Arrows indicate distorted gemma cups. Bars = 2 cm. (E–H) Top view of a 2-week-old thalli grown from gemmae. Bars = 5 mm. (I–L) Side view of E–H, respectively. Bars = 2 mm. (M–P) Enlarged images of rhizoids in I–L. Bars = 500 µm. **(Q–T)** Representative images used for rhizoid length quantification are shown in X. Five-day-old gemmalings of wild type, Mp*ropgap^ge^-1*, Mp*ren^ge^-1*, and Mp*ropgap^ge^* Mp*ren^ge^-2,* were grown on the upside-down medium. Bars = 1 mm. **(U)** Box-and-dot plots of the size of mature gemmae within the gemma cup formed on 3-week-old thalli. Different letters indicate a significant difference (one-way ANOVA; Tukey’s HSD, P <0.01). n = 50. **(V)** Gemmaling growth for 6 days. Asterisks indicate significant differences between mutant versus wild-type plants (Dunnett’s test). Bars = SE. n = 18. **(W)** Box-and-dot plots of flesh weight of 15-day-old gemmalings. Different letters indicate a significant difference (one-way ANOVA; Tukey’s HSD, P <0.05). n = 18. **(X)** Quantitative analysis of rhizoid elongation of gemmalings as shown in Q–T. Box-and-dot plots of the maximum length of rhizoids. Different letters indicate a significant difference (one-way ANOVA and Tukey’s HSD, P <0.01). n = 25–26.

Furthermore, both the Mp*ren^ge^* mutant and Mp*ropgap^ge^*Mp*ren^ge^* double mutants showed abnormalities in rhizoids, with the Mp*ren^ge^* mutant rhizoids being short and twisted and the Mp*ropgap^ge^* Mp*ren^ge^* double mutant rhizoids being thick and short (Figures 2M–T and 2X). All of these phenotypes were observed in the corresponding mutant alleles (Figure S5). These results suggest that MpREN predominantly regulates thallus growth and rhizoid elongation. Moreover, although the Mp*ropgap^ge^*single mutant exhibited a phenotype identical to that of the wild-type plants, the additive phenotype in the Mp*ropgap^ge^* Mp*ren^ge^* double mutants implied that MpROPGAP redundantly contributed to these regulatory processes with MpREN.

### MpREN and MpROPGAP play essential roles in gemma development

Given the abnormalities observed in gemma size in the single mutants of Mp*REN* and double mutants of Mp*ROPGAP* and Mp*REN* (Figure 2U), we performed a more detailed observation of gemma morphology. The wild-type gemmae had a flattened structure with two apical notches and dispersed oil bodies, predominantly in the margins (Figures 3A, F, and P; Kanazawa et al. 2022). The gemmae of the single Mp*ropgap^ge^* mutant exhibited a flattened structure with two apical notches, similar to those of the wild-type gemmae (Figures 3B, G and P). In contrast, the gemmae of the Mp*ren^ge^* single mutants were slightly smaller, had a rounded appearance, and had asymmetric structures with three or more notches (Figures 3C, H, and P). The gemmae of the Mp*ropgap^ge^* Mp*ren^ge^* double mutant were globular, and notch structures were unrecognized (Figures 3D, E, I, J, and P). Furthermore, a few gemmae of the double mutant had elongated rhizoids, even within the gemma cup (Figure 3E; arrowhead). To identify meristematic regions with active cell division in the gemmae, EdU staining was performed on 1-day-old gemmalings of mutant and wild-type plants. In the Mp*ropgap^ge^*single mutant, EdU-positive cells were observed mostly around the two apical notches, similar to the wild-type gemmae (Figures 3F, G, K, and L). In Mp*ren^ge^* single mutants, EdU-positive cells were detected around opposing notch structures and around ectopic notch structures (Figures 3H, M). In contrast, in the Mp*ropgap^ge^*Mp*ren^ge^* double mutants, EdU-positive cells were randomly distributed in a patchy pattern (Figures 3I, J, N, O, Q, and R). Abnormal directional growth of thalli was observed in the Mp*ren^ge^* single mutant and Mp*ropgap^ge^* Mp*ren^ge^* double mutant gemmae (Figures 2C, D, G, and H), which was attributed to the abnormal organization of meristematic regions during gemma development. These results suggest that MpREN and MpROPGAP are essential for appropriate patterning during gemma development.

**Figure 3.**
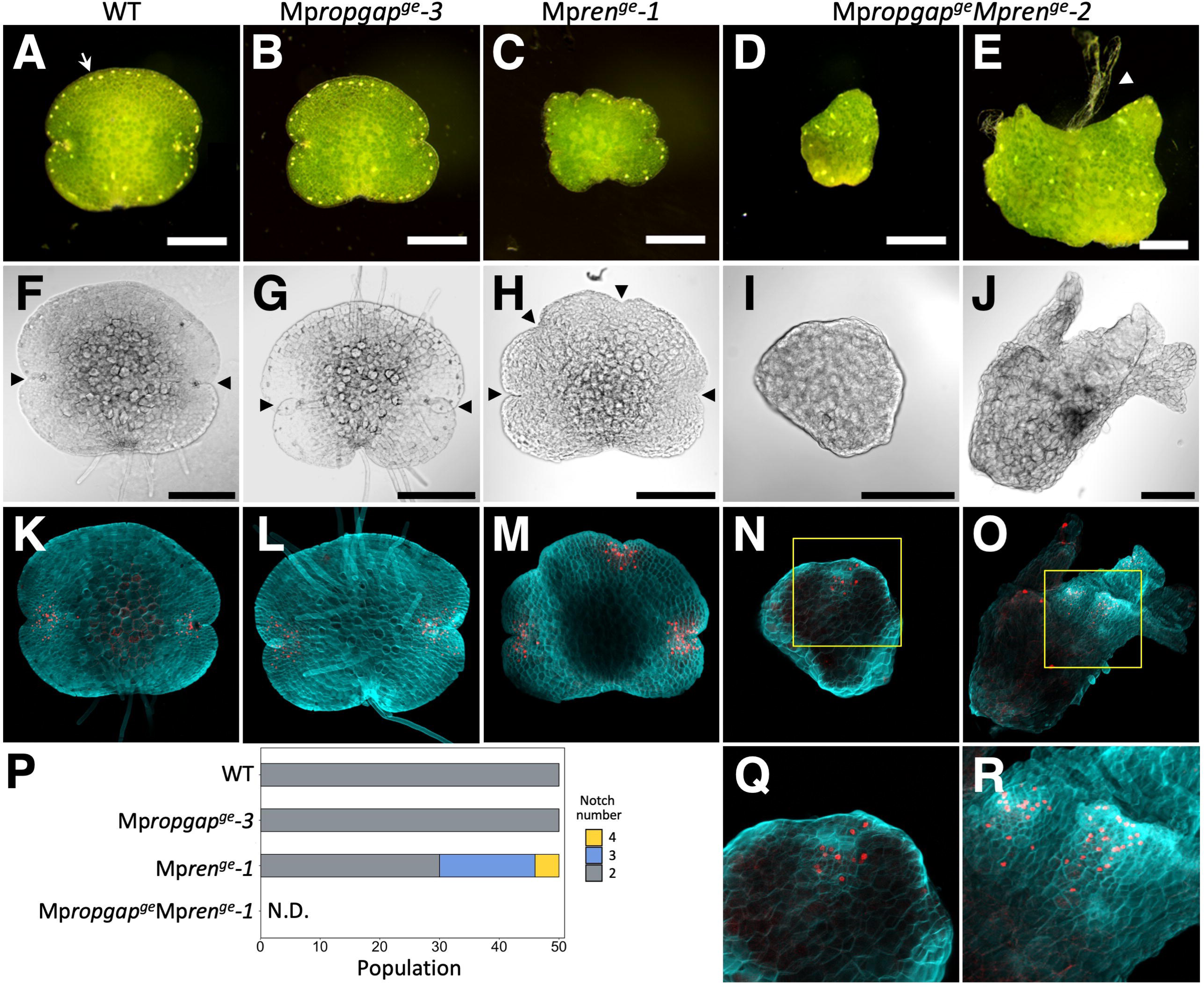
Morphology of gemmae in Mp*ROPGAP* and Mp*REN* loss-of-function mutants. Images of gemma in the wild-type **(A, F, and K),** Mp*ropgap^ge^***(B, G, and L),** Mp*ren^ge^* **(C, H, and M),** and Mp*ropgap^ge^* Mp*ren^ge^* **(D, E, I, J, N, and O)** lines. **(A–E)** Representative mature gemma. The white arrow indicates an oil body dispersed predominantly in the marginal area. The white arrowhead indicates elongated rhizoids. Bars = 250 µm. **(F–J)** Transmitted images and (**K–O, Q, and R)** z-projections of confocal images of gemmae labeled with EdU Alexa 555 (red signals) and SCRI Renaissance Stain 2200 (cyan signals). Arrowheads indicate the recognized notch structures. Enlarged images of the yellow squares in N and O are shown in Q and R, respectively. Bars = 250 µm. **(P)** Number of notch structures in the mature gemma within the gemma cup formed on 3-week-old thalli. n = 50.

### Significant roles of MpREN and MpROPGAP in the air-chamber development

In *M. polymorpha*, the dorsal side of the thallus is covered with photosynthetic organs called air chambers (Shimamura, 2016). Each air chamber had a single air pore distributed in a distinctive pattern on the dorsal surface of the thallus, as observed in the wild-type (Figures 4A, E) and Mp*ropgap^ge^* plants (Figures 4B, F). In the Mp*ropgap^ge^* mutant, a single-layered roof covering the entire air-chamber unit, equipped with a characteristic air pore composed of four layers with four cells and internal assimilatory filaments on the basal cells, was formed, as observed in the wild type (Figures 4I, J). However, in the Mp*ren^ge^* single mutant and Mp*ropgap^ge^* Mp*ren^ge^* double mutant, air pores were absent on the dorsal surface of the thallus, and the thallus epidermis ruptured (Figures 4C, D, G, H). In the Mp*ren^ge^* single mutants, the individual unit structure of the air chamber was ambiguous. From cross-sectional observations, assimilatory filaments were observed, air pores were absent, and the single-layered roof was partially lacking in the Mp*ren^ge^* single mutant (Figures 4C, G, K). In the Mp*ropgap^ge^* Mp*ren^ge^* double mutant, the phenotype was more severe, with the roof structure almost absent, exposing the assimilatory filaments (Figures 4D, H, and L). Similar abnormalities in air chamber formation have been observed in loss-of-function mutants of MpROP, where abnormal cell division patterns of dorsal epidermal cells around the apical notch lead to anomalies in the air chamber roof structure and air pore formation (Mulvey and Dolan 2023). Observation of the dorsal epidermis around the apical notch in 6-day-old gemmalings showed impairments in the proliferation of roof cells and air pore formation adjacent to the intercellular space in both the Mp*ren^ge^* single mutant and the Mp*ropgap^ge^* Mp*ren^ge^* double mutant (Figures 4M–P). These results revealed that MpREN and MpROPGAP redundantly regulate air chamber development by controlling the cell division pattern of epidermal cells in the meristematic region.

**Figure 4.**
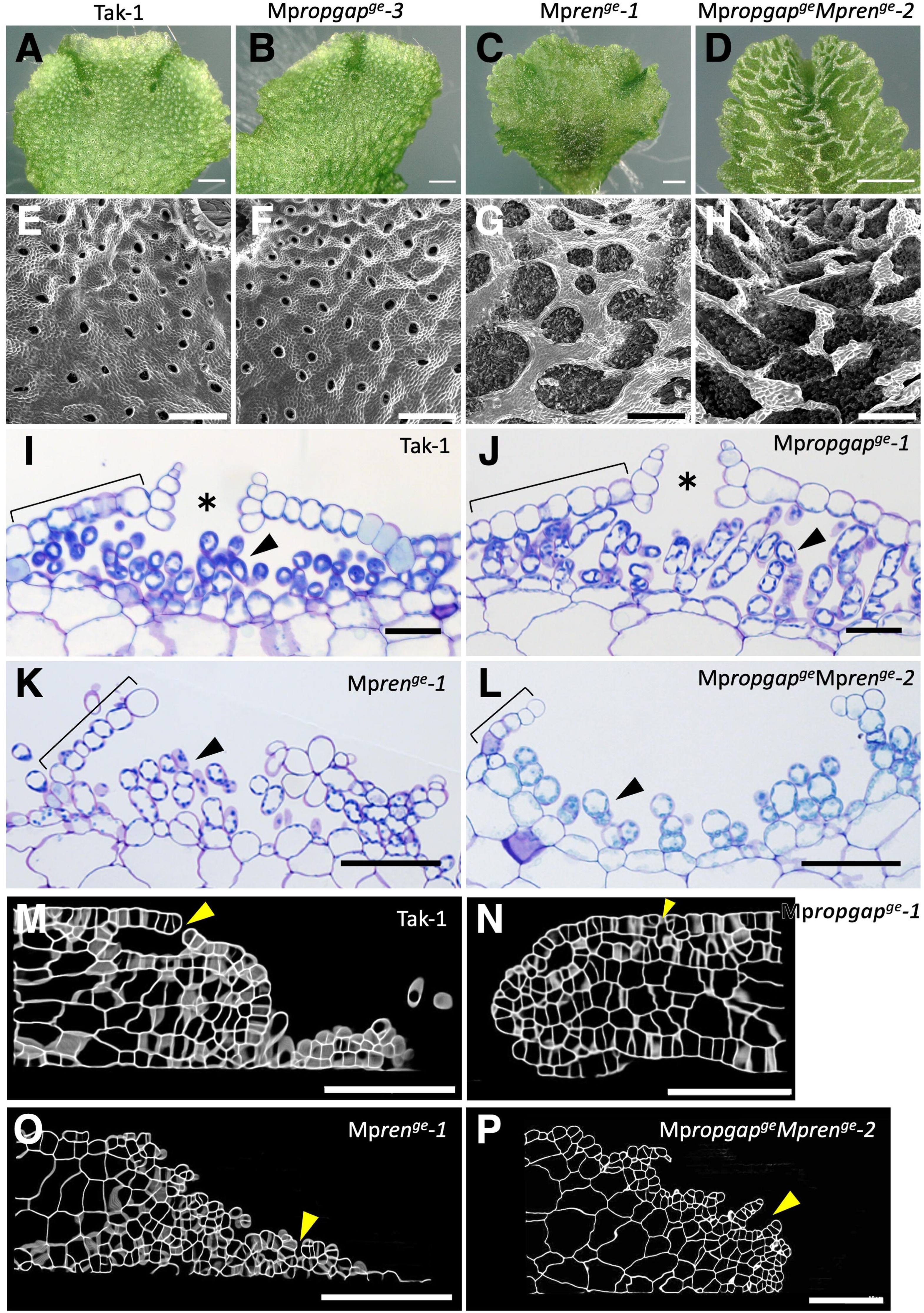
Morphology of the air chambers in Mp*ROPGAP* and Mp*REN* loss-of-function mutants. **(A–D)** Top view of a single thallus lobe in the 2-week-old gemmalings of wild type (A), Mp*ropgap^ge^-3* (B), Mp*ren^ge^-1* (C), and Mp*ropgap^ge^*Mp*ren^ge^-2* (D). Bars = 1 mm. **(E–H)** Scanning electron microscopic images of the dorsal surface of 2-week-old gemmalings. Bars = 500 µm. **(I–L)** Toluidine-blue-stained transverse sections of the mature air chambers. Brackets indicate an air-chamber roof. Asterisks indicate the air pores. Arrowheads indicate assimilatory filaments. Bars = 200 µm. **(M–P)** Optical cross-sections of developing air chambers near the meristematic notches in 6-day-old gemmalings. Images were processed in MorphoGraphX. Yellow arrowheads indicate the intercellular spaces. Bars = 100 µm.

### Proper regulation of the ROP activation/inactivation switching is essential for the development of air chambers

MpROPGAP and MpREN possess highly conserved GAP domains, which are predicted to activate the GTPase activity in ROP and facilitate the conversion of ROP from the active GTP-bound form to the inactive GDP-bound form. Consequently, we hypothesized that the levels of active ROP would increase in loss-of-function mutants of Mp*ROPGAP* and Mp*REN.* Conditional overexpression of *constitutively active* Mp*ROP* (Mp*ROP^CA^*) using an estradiol-inducible artificial promoter (*_pro_*Mp*E2F:XVE>>*) resulted in defects in air-chamber development similar to Mp*ren^ge^* single mutants and Mp*ropgap^ge^* Mp*ren^ge^* double mutants (Figure 5A). Furthermore, conditional overexpression of MpROPGEF (KARAPPO) led to a similar air-chamber phenotype (Figure 5A). These results strongly suggest that the phenotypes of air-chamber development both in Mp*ren^ge^* single mutants and Mp*ropgap^ge^* Mp*ren^ge^* double mutants are associated with defects in the promotion of GTP hydrolysis by MpROP.

**Figure 5.**
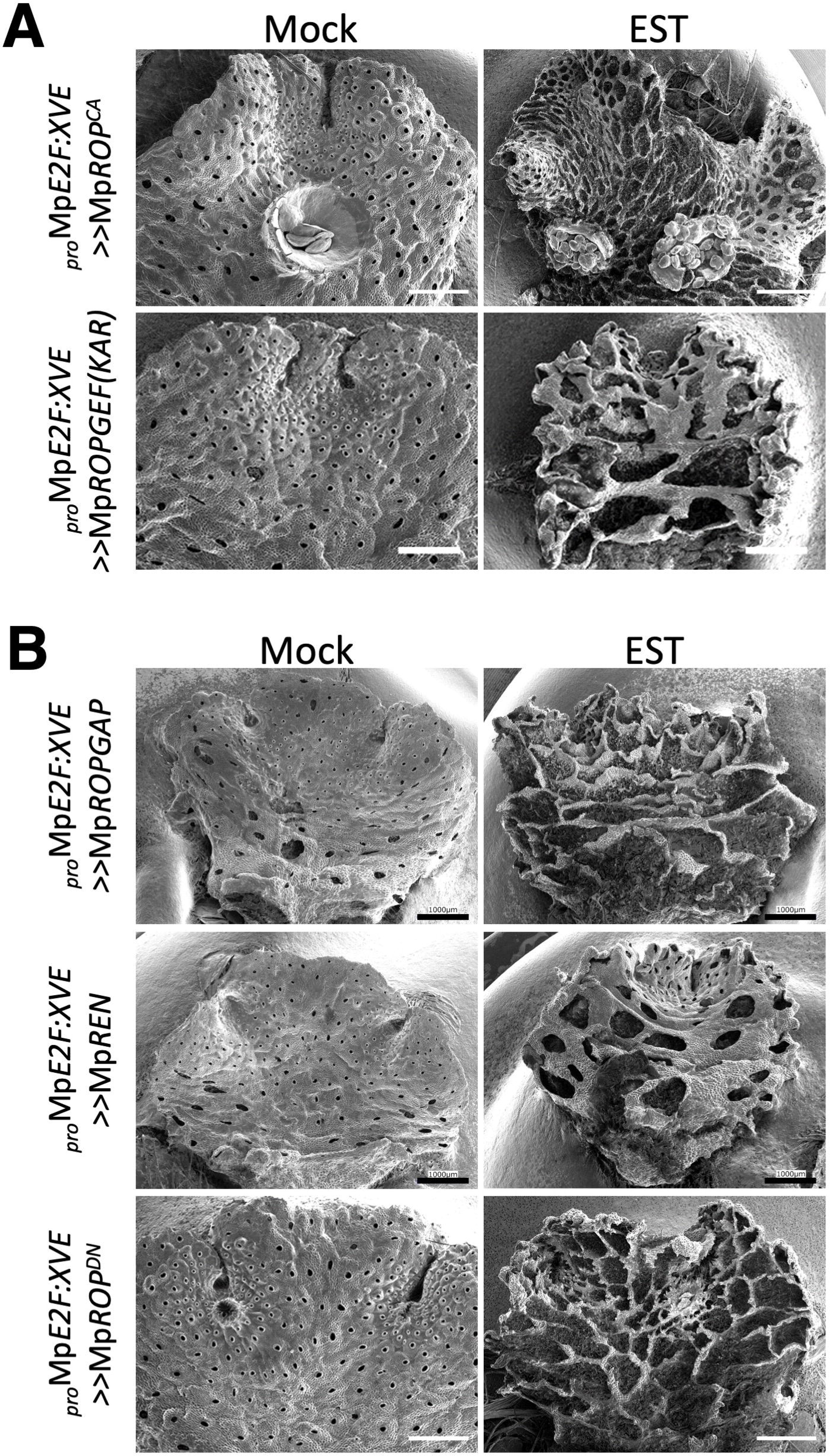
Air-chamber impairments induced by an excess of active or inactive MpROP. Scanning electron microscopic images of the dorsal surface of a 2-week-old thalli of **(A)** *_pro_*Mp*E2F:XVE>>*Mp*ROP^CA^*and *_pro_*Mp*E2F:XVE>>*Mp*ROPGEF(KAR)* and **(B)** *_pro_*Mp*E2F:XVE>>*Mp*ROPGAP*, *_pro_*Mp*E2F:XVE>>*Mp*REN*, and *_pro_*Mp*E2F:XVE>>*Mp*ROP^DN^*, grown in the presence (EST) or absence (Mock) of 10 µM LJ-estradiol. Bars = 1 mm.

On the other hand, conditional overexpression of Mp*ROPGAP* and Mp*REN* also induced air-pore defects and dorsal epidermal rupture, respectively (Figure 5B). Additionally, the conditional overexpression of *dominant-negative* Mp*ROP* (Mp*ROP^DN^*) led to a similar phenotype (Figure 5B). These results suggest that an increase in inactive MpROP (a decrease in active MpROP) impairs air chamber formation, implying the essential role of proper switching/cycling of MpROP activation and inactivation in the process of air chamber development.

In the genome of *M. polymorpha*, single copies of PRONE-type ROPGEF and DHR-type SPIKE (MpSPK) were predicted to be the GEFs that activate MpROP. Although MpROPGEF (KAR) is essential for the initiation of gemmae from the initial gemma cells on the basal floor of the gemma cup, it has no significant effect on air chamber development (Hiwatashi et al. 2019). Therefore, to investigate the role of MpSPK, we generated transgenic plants expressing an artificial microRNA (amiRNA) targeting Mp*SPK* mRNA under the control of the estradiol-inducible artificial transcription factor XVE (*_pro_*Mp*E2F:XVE>>amiR_*Mp*SPK^Mpmir160^*; Figures S6A, B). Conditional suppression of Mp*SPK* resulted in pronounced inhibition of thallus growth and defects in air chamber formation (Figure 6). Consistent results were obtained for the different knock-downed Mp*SPK* lines (Figure S6C). These results suggested that MpSPK plays a crucial role in air chamber development. MpROPGEF and MpSPK are responsible for MpROP activation during organogenesis in *M. polymorpha*.

**Figure 6.**
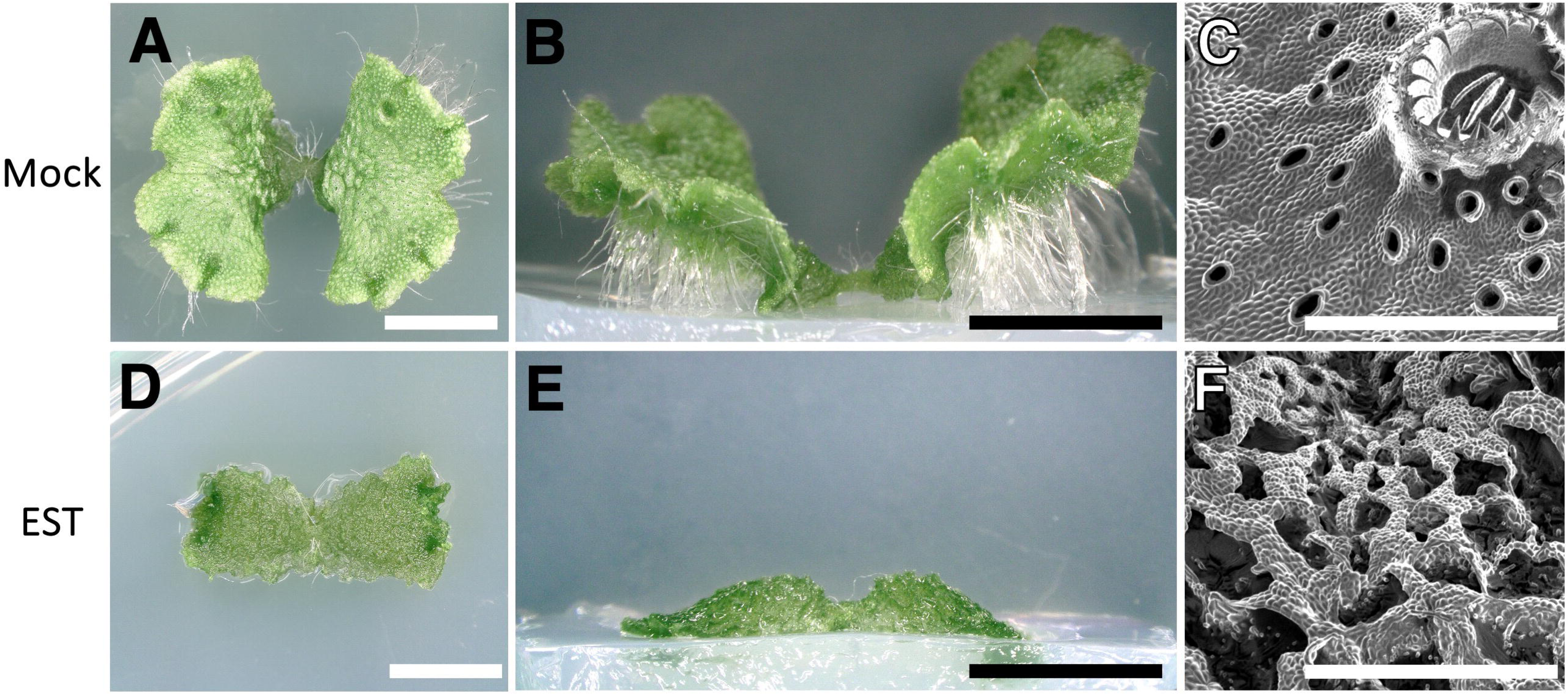
Air-chamber defects by conditional suppression of Mp*SPK*. Top view **(A and D),** side view **(B and E)**, and dorsal surface **(C and F)** of 2-week-old thalli (*_pro_*Mp*E2F:XVE>>amiR1_*Mp*SPK*^Mp*mir160*^*-1*) grown in the absence (Mock; A–C) or presence (EST; D–F) of 10 µM LJ-estradiol. Scale bar = 5 mm (A, B, D, and E). Scale bars = 1 mm (C and F).

## Discussion

### RhoGAPs are necessary for the morphogenesis of *M. polymorpha*

*M. polymorpha* possesses single-copy genes encoding ROPGAP (CRIB-GAP) and REN (PH-GAP), which harbor a GAP domain predicted to promote the GTPase activity of ROP. These genes were expressed throughout the life cycle of *M. polymorpha* (Figures 1 and S3). A previous report showed that the Mp*REN* mutant exhibited a curly rhizoid phenotype, suggesting the involvement of MpREN in rhizoid elongation (Honkanen et al. 2016). In this study, loss-of-function mutants of Mp*REN* showed systemic organogenetic abnormalities, including air chambers, gemma cups, and rhizoids, accompanied by significant inhibition of thallus growth and abnormal gemma morphology (Figures 2, 3, and 4). These findings highlighted the crucial regulatory roles of MpREN in various developmental processes. In contrast, the single mutant of Mp*ROPGAP* displayed no apparent defects in organ formation or thallus growth (Figures 2, 3, and 4), suggesting relatively minor roles for MpROPGAP in these processes. However, the Mp*ropgap^ge^* Mp*ren^ge^* double mutant exhibited additive and more severe defects in these developmental processes than the Mp*ren^ge^* single mutant (Figures 2U–X). Despite the different domain structures of MpROPGAP and MpREN, except for the GAP domain (Figures S1 and S2), these data suggest that MpREN and MpROPGAP function redundantly in various developmental processes. The developmental processes regulated by MpREN and MpROPGAP are diverse, such as tip growth in single cells observed in rhizoid elongation, two-dimensional proliferation of protodermal cells in air-chamber roof expansion, and three-dimensional patterning in gemma development. Considering that the developmental processes regulated by MpREN and MpROPGAP align well with those regulated by MpROP (Rong et al. 2022; Mulvey and Dolan 2023), MpROP inactivation by these two RhoGAPs is essential for these processes.

In the angiosperm *A. thaliana*, RhoGAPs contribute to cell polarity by restricting the intracellular localization of ROP proteins and creating anisotropic active ROP domains (Feiguelman et al. 2018). During metaxylem development, it has been experimentally demonstrated that the simultaneous expression of ROP, ROPGEF, and ROPGAP results in the formation of active ROP domains, suggesting that the self-organization properties of the ROP-ROPGEF-ROPGAP modules result in cellular patterning (Oda and Fukuda 2012). In the pavement cells of *A. thaliana*, the restriction of active ROP domains to multiple locations allows for jigsaw-like cell morphogenesis (Lauster et al. 2022; Zhang et al. 2022). However, in tip-growing cells, such as the root hair and pollen tube, ROPGAPs and RENs inactivate ROP by localizing to the subapical and apical domains, thereby spatiotemporally limiting the active ROP domain within the apical part of the cell (Ou and Yi 2022). Since the loss of function of RhoGAPs and ROP alone caused tip growth inhibition in the rhizoids, it is expected that a similar regulatory mechanism of cell polarity formation may underlie cellular morphogenesis in *M. polymorpha*.

The importance of ROP signaling has mainly been shown in cell polarity formation in various cellular morphogenesis processes, as mentioned above; however, recently, it has become important in cell division orientation (Müller 2023). In *Arabidopsis* early embryogenesis and root meristem, the inactivation of ROP in the cortical division zone/site (CDZ/CDS) affects the cytoskeletal organization in the CDZ/CDS, allowing cell division orientation. AtREN1 and AtREN2 are responsible for ROP inactivation at the CDZ/CDS (Stöckle et al. 2016). In *M. polymorpha*, loss-of-function mutants of Mp*ROP* failed to form an air chamber containing a roof and air pores (Mulvey and Dolan 2023). This is attributed to the loss of control over the orientation of cell division, especially in the protodermal cell layers near the meristem notch (Rong et al. 2022; Mulvey and Dolan 2023). In this study, we demonstrated that the loss of function of Mp*REN* led to an abnormal orientation of cell division in the protodermal cell layers and the manifestation of an incomplete phenotype in air-chamber roofs equipped with air pores (Figure 4 K, O). This suggests a primary and essential role for MpREN in the orientation of cell division planes during single-layered roof cell proliferation. Further investigation of the subcellular localization of MpREN and the effects of MpREN mutations on the localized activation of MpROP in the cell may provide insights into the common mechanisms between bryophytes and angiosperms. The role of the ROP signaling pathway in the control of cell division orientation is fundamental and conserved across land plants.

### Regulation of ROP cycling is essential for air chamber development

The histological process of air chamber development on the dorsal surface of the thallus in *M. polymorpha* has been well described. Initially, dorsal epidermal cells near the meristem notch undergo periclinal cell division, giving rise to protodermal and sub-protodermal cells (Ishizaki 2015). Following this division, the intercellular space becomes apparent as an initial aperture between the anticlinal walls of the protodermal cells, manifesting as a gap at the junction where the four protodermal cells meet. The base of the initial aperture widened, and the primary air chamber developed through successive anticlinal cell divisions and the growth of both protodermal and sub-protodermal cells surrounding the intercellular space. The protodermal cell layers form the air chamber roof via anticlinal cell division, whereas the sub-protodermal cell layer forms the air chamber floor through a similar division. In the four protodermal cells surrounding the intercellular gap of the roof, oblique and periclinal divisions lead to the formation of a four-layered air-pore structure (Mulvey and Dolan 2023). The loss of MpROP function disrupts the formation of both the air-chamber roof and air pores, indicating a crucial role for MpROP in the regulation of these processes (Mulvey and Dolan 2023). Our study revealed that the loss of MpREN function (Figure 4) and the overexpression of the constitutively active form of ROP (MpROP^CA^) and MpROPGEF resulted in abnormalities in the air-chamber roof and air pore formation, suggesting that the presence of excess active ROP impaired air-chamber organogenesis (Figure 5A). Interestingly, the overexpression of MpROPGAP or MpREN and dominant-negative mutated ROP (MpROP^DN^) also led to abnormalities in the air-chamber roof and air pore formation, indicating that an increase in the inactive form of ROP (a decrease in the active form of ROP) impaired organ formation (Figure 5B). These results emphasize the importance of proper control of the ROP active/inactive status through ROP cycling during air chamber development.

Our results showed that MpREN plays a predominant role in ROP inactivation during air chamber morphogenesis. However, the factors responsible for ROP activation during air chamber morphogenesis remain unknown. Among the candidate GEFs in *M. polymorpha*, the loss-of-function mutation of the PRONE-type GEF, MpROPGEF (KAR), exhibited no detectable phenotype during air chamber development, indicating that MpROPGEF may not be a major player in this process (Hiwatashi et al. 2019). In this study, we demonstrated that post-transcriptional suppression of the DOCK family GEF Mp*SPK* leads to severe air chamber defects (Figure 6), indicating the crucial role of MpSPK in MpROP activation during air chamber development. Although ROP in *M. polymorpha* regulates various developmental and organ-forming processes, the genome lacks homologous genes encoding known effectors in angiosperms, and the factors regulating diverse downstream signal transduction pathways remain to be elucidated. Functional differentiation between the two types of GEFs, as demonstrated in this study and a previous study (Hiwatashi et al. 2019), may be achieved by different compositions of protein complexes and their subcellular localization, which mediates MpROP signaling. Addressing the subcellular localization of MpROPGEF and MpSPK together with MpROP-activated domains and identifying their unknown interacting factors will shed light on the fundamental mechanism of diverse MpROP functions with a minimum set of regulatory components in *M. polymorpha*.

## Material and methods

### Phylogenetic analysis

To search for MpROPGAP and MpREN, the amino acid sequences of *M. polymorpha* and *P. patens* were collected from MarpolBase (http://marchantia.info/, Bowman et al. 2017) and *A. thaliana* from TAIR (http://www.arabidopsis.org). Multiple alignment diagrams were generated using MUSCLE (Edgar 2004) in MEGA7 (Kumar et al. 2016), a phylogenetic analysis integration software package. The generated multiple alignment diagrams were processed using BoxShade (https://embnet.vital-it.ch/software/BOX_form.html).

A homology search (BLASTP) was performed on the amino acid sequences of the proteins encoded in the genome, using the MpROPGAP and MpREN sequences as queries. Among the sequences used, *M. polymorpha* (Mp6g11120, MpROPGAP; Mp8g09680, MpREN), *Klebsormidium nitens* (kfl00008_0150; kfl00251_0100), *Physcomitrium patens* (Pp3c26_5960, Pp3c26_4490, Pp3c4_24980, Pp3c4_16800, Pp3c 13_4010, Pp3c3_5940; Pp3c9_17460), *Amborella trichopoda* (AmTr_scaffold00027.1, AmTr_scaffold 00040.52, AmTr_scaffold00057.110; AmTr_scaffold00135.11, AmTr_scaffold00063.78), and *Selaginella moellendorffii* (Sm_13019, Sm_13258; Sm_442380) were obtained from MarpolBase (http://marchantia.info/), *Arabidopsis thaliana* (AT2G46710, AT4G03100, AT1G08340, AT3G11490, AT5G22400; AT4G24580, AT5G12150, AT5G19390) were from Phytozome (https://phytozome.jgi. Doe.gov/pv/portal.html), *Azolla filiculoides* (Azfi_s0055.g033926, Azfi_s0108.g045281, Azfi_s0037.g026056, Azfi_s1557.g104316, Azfi_s0037.g025995, Azfi_s0001.g000579, Azfi_s0081.g 038659; Azfi_s0173.g055873, Azfi_s0028.g023819), and *Salvinia 13ucullate* (Sacu_s0040.g012358, Sacu_s0074.g017279, Sacu_s0010.g004927, Sacu_s0004.g002103, Sacu_s0014.g006099, Sacu_s0013.g005777, Sacu_s0013.0013.g005777, Sacu_s0010.g004920, Sacu_s0022.g008570, and Sacu_s0056.g014542) were obtained from FernBase (https://fernbase.org/), *Picea abies* (MA_10436113g0010) from Congenie (https://congenie.org/), and *Anthoceros angustus* (AANG004126, AANG013186) from Zhang et al. (2020). Each sequence was multiple-aligned using MEGAⅩ’s MUSCLE program (Edgar 2004), and only the sequence portions of the major domain structures were used. Phylogenetic analysis was performed using PhyML (Guindon et al. 2010) with the Maximum Likelihood method, with bootstrap values of 100 for ROPGAP using the LG+G model and REN using the JTT+G model.

### Plant materials and growth conditions

*M. polymorpha* accession Takaragaike-1 (Tak-1; Ishizaki et al. 2008) was used as the wild type. Supplementary Table 1 lists the plants used in this study. Growth conditions have been described previously (Ishizaki et al. 2008). Plants were cultured on 1/2-strength Gamborg’s B5 medium (Gamborg et al. 1968) containing 1.0% (w/v) agar at 22°C under continuous white light.

### Quantitative RT-PCR analysis

Total RNA (180 ng of total RNA) was reverse-transcribed in a 10 µL reaction mixture using ReverTra Ace qPCR RT Master Mix with gDNA remover (TOYOBO). After the reaction, cDNA samples were diluted with 90 µL of distilled water. Templates were amplified in KOD SYBR qRT-PCR Mix (TOYOBO) using a Light Cycler Nano Real-time PCR Detection System (Roche Applied Science). The PCR was performed according to the manufacturer’s instructions. The primer pairs used in the experiments are listed in Table S1. Mp*EF1*_Cl_ (Mp*3g23400*) was used as an internal control.

### Generation of transformants for promoter–reporter analysis

To construct *_pro_*Mp*ROPGAP:GUS* and *_pro_*Mp*REN:GUS*, a 3837 bp upstream genomic region from the start codon of Mp*ROPGAP* (Mp*ROPGAPpro*) and a 5045 bp upstream genomic region from the start codon of Mp*REN* (*_pro_*Mp*REN*) were amplified from wild-type Tak-1 genomic DNA using PCR with the appropriate primer pairs (Table S1) and cloned into the pENTR/D-TOPO cloning vector (Life Technologies). These entry vectors were used in the Gateway LR reaction (Life Technologies) with the Gateway binary vector pMpGWB104 (Ishizaki et al. 2015) to generate *_pro_*Mp*ROPGAP:GUS* and *_pro_*Mp*REN:GUS* binary constructs. The *_pro_*Mp*ROPGAP:GUS* and *_pro_*Mp*REN:GUS* binary vectors were introduced into Tak-1 thalli by Agrobacterium-mediated transformation as previously described (Kubota et al. 2013). The transformants were selected using 10 mg/mL hygromycin B and 100 mg/mL cefotaxime.

### CRISPR/Cas9-based genome editing of MpROPGAP and MpREN

Loss-of-function mutants of Mp*ROPGAP* and Mp*REN* were generated using the CRISPR/Cas9 system as described previously (Sugano et al. 2014; Sugano et al. 2018). According to the guidelines of the CRISPRdirect website (https://crispr.dbcls.jp), we selected two target sequences for Mp*ROPGAP*, one located in the 2^nd^ exon and the other located in the 3^rd^ exon of Mp*ROPGAP*, whereas three target sequences for Mp*REN* were located in the 4^th^ exon, 8^th^ exon, and 11^th^ exons of Mp*REN* (Figure S2). Synthetic oligo DNAs for the respective target sites shown in Table S1 were annealed, inserted into the entry vector pMpGE_En03 (GenBank Accession #LC090755), and introduced into the destination vectors pMpGE010 or pMpGE011 (Sugano et al. 2014; Sugano et al. 2018). The vectors were introduced into the regenerating thalli of Tak-1 via A*. tumefaciens* GV2260 (Kubota et al. 2013), and transformants were selected with 10 mg/mL hygromycin B or 0.5 mM chlorsulfuron and 100 mg/mL cefotaxime. Genomic DNA was isolated from the transformants and amplified from the target region using PCR. The PCR product was used to sequence the respective target sites using an ABI 3100 Genetic Analyzer (Applied Biosystems). Mutants were named according to the nomenclatural rules for *M. polymorpha* (Bowman et al. 2016).

### Histology and light microscopy

For histochemical GUS staining, *_pro_*Mp*ROPGAP:GUS* and *_pro_*Mp*REN:GUS* transgenic plants were grown on a half-strength B5 medium containing 1% agar for different periods under continuous white light irradiation. GUS staining was performed as previously described, and at least three independent lines were observed for GUS staining patterns using an M205 FA stereoscopic microscope (Leica Microsystems) equipped with a CCD camera (DFC7000 T; Leica Microsystems). For systemic phenotypic analysis, top and side views of the plants were captured using a VHX-5000 digital microscope (KEYENCE). For plastic-embedded sectioning, 2-week-old thalli developed from gemmae were dissected into small pieces and transferred to a fixative solution (2% glutaraldehyde in 0.1 M HEPES buffer, pH 7.2), evacuated with a water aspirator until the specimens sank, and fixed for 3 days at room temperature (22–25°C). The samples were dehydrated using a graded ethanol series and embedded in Technovit 7100 plastic resin. Semi-thin sections (3 µm thickness) were obtained using a microtome (HM 335E; Leica Microsystems) and stained with 0.2% toluidine blue O for observation under a BX51 light microscope (OLYMPUS). For scanning electron microscopy, plant samples were frozen in liquid nitrogen and directly observed using a VHX-D510 microscope (KEYENCE).

### Quantitative analysis of gemmaling growth and rhizoid elongation

To quantify the growth of each genotype, gemmae were incubated on 1/2-strength B5 1% agar medium for 0, 2, 4, and 6 days and imaged using an M205 FA stereoscopic microscope (Leica) equipped with a CCD camera (DFC7000 T; Leica). The area of individual plants in the images was measured using the ImageJ Fiji software (Schindelin et al. 2012). Dunnett’s test, followed by a one-way ANOVA, was performed for statistical analysis. To analyze gemma morphology, the size and number of notch structures were measured in gemmae from mature gemma cups of each genotype. Images were captured with an M205 FA stereoscopic microscope (Leica) equipped with a CCD camera (DFC7000 T; Leica) and analyzed using ImageJ Fiji. To measure fresh weight, 18 gemmae were cultured on 1/2-strength B5 medium for 15 d, and individuals were weighed. To measure the maximum rhizoid length, 25–26 gemmae were grown upside down on 1/2-strength B5 medium for 5 days, and images were captured using an M205 FA stereoscopic microscope (Leica) equipped with a CCD camera (DFC7000 T; Leica). The longest rhizoid, defined as the vertical distance from the agar gel surface, was measured for each gemmalings by ImageJ Fiji. Tukey-Kramer’s test following one-way ANOVA was performed for statistical analysis.

### Confocal imaging of the initial stage of the air chamber development

For histochemical observation of the apical notch, 5-day-old gemallings were fixed in 4% (w/v) formaldehyde in 1×phosphate-buffered saline and cleared with ClearSee_a_ (Kurihara et al. 2021), as described in Mulvey and Dolan (2023). The cell walls of specimens were stained with 0.2% (v/v) Renaissance SR2200 in ClearSee_a_ as previously described (Mulvey and Dolan 2023). The confocal images were obtained with a confocal laser-scanning microscope (FV1000; Olympus) equipped with 405 nm LD laser lines. Serial images were acquired at 0.5-µm intervals in depth with a 30 × 1.05 N.A. oil immersion objective (UPLSAPO30XS; Olympus) using silicon immersion oil. The z-stacked images were processed to obtain vertical optical sections using the MorphoGraphX program (Barbier De Reuille et al. 2015; Vijayan et al. 2021).

### Generation of transgenic plants overexpressing ROP-related genes

To generate *_pro_*Mp*E2F:XVE>>MpROPGAP* and *_pro_*Mp*E2F:XVE>>MpREN*, the Mp*ROPGAP* and Mp*REN* coding sequences were amplified using PCR by KOD Plus Neo DNA polymerase (TOYOBO) with the appropriate primer pairs (Table S1) and cloned into the pENTR/D-TOPO cloning vector (Life Technologies). To generate *_pro_*Mp*E2F:XVE>>MpROP^CA^, _pro_*Mp*E2F:XVE>>MpROP^DN^,* and *_pro_*Mp*E2F:XVE>>MpROPGEF(KAR),* we used the coding sequences previously cloned in the pENTR/D-TOPO cloning vector by Hiwatashi et al. (2019). The entry vectors were used in the Gateway LR reaction with the Gateway binary vectors pMpGWB168 and pMpGWB368 (Ishida et al. 2022). These binary vectors were introduced into the regenerating thalli of Tak-1 as previously described (Kubota et al. 2013). Transformants were selected with 10 mg/mL hygromycin B or 0.5 mM chlorsulfuron and 100 mg/mL cefotaxime. Transformants were named according to the nomenclatural rules for *M. polymorpha* (Bowman et al. 2016).

### Conditional knockdown of Mp*SPK*

To generate *_pro_*Mp*E2F:XVE>>amiR_MpSPK^Mpmir160^*, we designed artificial-micro RNAs (amiRs) with an Mp*MIR160* backbone targeting the Mp*SPK* transcript (Figures S6A and S6B), according to Flores-Sandoval et al. (2016). We annealed a pair of synthesized 85-mer oligonucleotides for the sequence spanning from the miR to the miR* (Figure S6) of *amiR_*Mp*SPK^Mpmir160^*(Table S1) and cloned by golden gate method into PaqCI cloning site of the vector pMpAmiR_160_En01, which contains the MpMIR160 backbone sequence between Gateway cassettes (attL1-attL2). Thereafter, *amiR_*Mp*SPK^Mpmir160^*sequences were cloned into the Gateway destination vector pMpGWB168 (Ishida et al. 2022) by the Gateway LR reaction. These binary vectors were introduced into the regenerating thalli of Tak-1 as previously described (Kubota et al. 2013). The transformants were selected using 10 mg/mL hygromycin B and 100 mg/mL cefotaxime.

### Phenotypic analysis of conditional overexpression and knockdown of genes

To evaluate the effects of conditional overexpression of genes, gemmae of these transgenic plants were cultured for two weeks on the 1/2-strength Gamborg’s B5 medium plates containing 10 μM β-estradiol (Fujifilm) or the equivalent volume (0.1% [v/v]) of DMSO (Fujifilm) as a mock control. The plant samples were frozen in liquid nitrogen and directly observed under a VHX-D510 microscope (KEYENCE, Osaka, Japan). For electron scanning microscopy, the plants were frozen in liquid nitrogen and directly observed under a VHX-D510 microscope (KEYENCE).

## Acknowledgments

The authors thank Eri Okada and Chiho Hirata for their technical assistance. We would like to thank Editage (www.editage.jp) for English language editing.

## Data Availability Statement

The authors confirm that the data supporting the findings of this study are available within the article and its supplementary materials.

